# Lymphocyte subpopulations and mast cells intestinal changes as indicators of inflammatory bowel disease in dogs

**DOI:** 10.1101/723536

**Authors:** Andrés Espinoza-Zambrano, Carlos Manuel González

**Affiliations:** Escuela de Medicina Veterinaria, Facultad de Ciencias de la Vida, Universidad Andres Bello, Santiago, Chile

**Keywords:** dog, inflammatory bowel disease, immunohistochemistry, Tlymphocytes, mast cells

## Abstract

Inflammatory bowel disease (IBD) is a disease with recurring gastrointestinal symptoms. Lymphocytes and mast cells are proposed as important components in the immunopathology of IBD in dogs. Mast cells depend on degranulation, a process that compromises mucosal permeability and normal intestinal barrier function, which alters the normal inflammatory process by allowing recruitment of lymphocytes in dogs with IBD. In this study, T and B lymphocyte populations and mast cells were examined in situ in 39 intestinal samples of dogs affected by IBD, by immunohistochemistry. Both T lymphocytes and mast cells numbers were significantly higher in the lamina propria of the intestinal wall of dogs with IBD compared with control dogs. Out of the total number of mast cells detected by CD117 expression significantly less cells appear to be granulated according to granule staining with Toluidine Blue, suggesting that an important degranulation process takes place in IBD. Single and double immune staining for tryptase and chymase showed that mast cells can express only one or both enzymes. Tryptase positive cells were significantly higher in number that chymase positive and tryptase/chymase positive cells. T lymphocytes were concentrated mostly at the upper portion of the intestinal villi lamina propria while mast cells were distributed mainly among crypts. These results suggest that populations of T lymphocytes and mast cells play a role in the immunopathology and development of IBD in dogs, also these changes could be helpful as complementary indicators of canine IBD.

## Introduction

Canine inflammatory bowel disease (IBD) is a group of gastrointestinal disorders characterized by persistent or recurring gastrointestinal symptoms with histologic evidence of inflammatory cell infiltration (1). There are different forms of IBD in dogs, depending on the main cell type involved and the affected portion of the intestine where the infiltration takes place, being lymphocytic-plasmacytic enteritis the most common form (2). The etiology of IBD remains unknown. Several studies suggest that these diseases result from inappropriate immune responses to the intestinal microbiome in genetically susceptible individuals (3–6). There are several factors involved including microbiome, environmental factors, genetic predisposition, and changes in the immune response of the individual, which may lead to loss of tolerance to the endogenous flora and the development of chronic inflammation of the gastrointestinal tract (7,8).

Different studies (9–12) suggest that a primary defect in the recognition of commensal bacteria or bacterial pathogens in the intestinal lumen, may lead to an increase in the production of IL-23, that induce naïve T cells to differentiate into T-helper (Th) lymphocytes, which release large amounts of proinflammatory cytokines (13,14). Theses cytokines damage the intestinal epithelium, allowing other pathogens to invade the lamina propria, driving more naïve T cell to differentiate into Th cells (4). Kleinschmidt and colleagues reported the increase of Th lymphocytes in IBD dogs, finding that lymphocytic-plasmacytic enteritis is mediated by Th1 lymphocytes, while eosinophilic gastroenteritis is mediated by hypersensitivity reactions modulated by Th2 lymphocytes, suggesting that there are different pathologic mechanisms (15). However, Heilmann and Suchodolski mentioned that in contrast to the results in human IBD, a distinct T-helper lymphocyte profile has not been clearly demonstrated to exist in canine or feline inflammatory bowel disease at this time (16). Overall, these studies evidence the importance of T lymphocytes in the adaptive immune system response during the development of IBD in dogs.

Besides T lymphocytes, B cells had been related to immunologic abnormalities in IBD in dogs too, but their number is usually lower compared with T lymphocytes (17). B cells perform several immunological functions but have been considered mainly as positive regulators of immune responses and central contributors to the pathogenesis of immune-related diseases because of their ability to produce antibodies, especially during the period between innate and adaptive immunity (18,19). The initial event that drives naïve cells to differentiate in B cells in IBD remains unknown, however, abrogation of the oral tolerance to commensal bacteria, lymphoid neogenesis, and hyperplasia in lesions may be involved (20).

Different molecular markers are used to determinate the populations of lymphocytes existing in the intestinal mucosa. The cluster of differentiation 3 (CD3) is a protein complex located on T lymphocyte cell surface and makes an ideal target for T lymphocyte in tissue sections. For detecting B cells, CD79a which is expressed by B lymphocytes during differentiation form early pre-B cell stage through to plasma cells, makes it a useful marker for mature B lymphocyte (21), or CD20, which is expressed in pre-, naïve and mature B-lymphocytes (22).

In addition to lymphocytes, mast cells and its mediators have also been associated with the pathogenesis of IBD. Mast cells are haematopoietic cells that arise from bone marrow and develop into mature mast cells under the influence of local growth factors, in particular, stem cell factor (SCF) (23). SCF, the ligand for the CD117/c-Kit receptor, is essential for mast cell proliferation, development, migration, survival and mediator release (24). Normally, mast cells can be found close to blood vessels or nerves beneath epithelial surfaces, where these cells are stimulated by environmental antigens (25). Within seconds of stimulation, mast cells can undergo degranulation, rapidly releasing pre-formed mediators present within cytoplasmic granules, including histamine, proteases, and tumor necrosis factor-alpha (TNF-a) (26). This affects the intestinal barrier function by increasing mucosal permeability. GI mast cell can contribute to an ongoing inflammatory process in the GI tract, allowing recruitment of granulocytes and lymphocytes to the site of the infection (27). Also, it had been speculated that interaction between commensal bacteria and mast cells affects the gastrointestinal barrier, promoting mucosal penetration of pathogens in the intestinal lamina propria perpetuating tissue damage and adaptive immune cells infiltration (28).

Commonly, toluidine blue staining of granules is the method used for the identification and quantification of mast cells in most tissues (29,30), but it is not always possible to detect fully degranulated mast cells with this technique (31). Alternately, CD117, which is a mast/stem cell grown factor receptor (SCFR) found in the membrane of mast cells, had been targeted to determinate activated mast cell in several tissues, including degranulated mast cells in the gastrointestinal mucosa (32,33).

Mast cells contain preformed granules that include heparin, histamine, cytokines, chemotactic factors, proteases and lipid derivatives that fulfill several biological functions, among which are phagocytosis, release of vasoactive substances and chemotaxis. (34). Mast cells are particularly rich in serine proteases stored and released from the secretory granules of mast cells. They present two subtypes, the mast cells, which contain only chymase and the mast cells that produce only tryptase and are located in the mucous membranes, close to T helper cells (Th2 type that release IL-4, 5, 6 and 13) (35), particularly in the intestinal lamina propria. These proteases promote vascular permeability through several mechanisms. They contribute to tissue remodeling through selective proteolysis of the matrix and the activation of metalloproteinases (36), in addition they promote the chemotaxis of inflammatory cells that surround the innate immune response (37). Changes in the number of mast cells in several anatomical sites and/or evidence of degranulation have been observed in a wide range of responses including hypersensitivity reactions (36,38), fibrosis, autoimmune pathology, inflammatory diseases, and neoplasia (39–41).

Lymphocytes and mast cells seem to play a role in the development and maintenance of IBD in dogs, considering they are an important cell component in lesions associated with chronic and autoimmune diseases. Changes in lymphocyte and mast cells populations could help in the diagnosis of IBD and contributes to improving understanding of this disease. Thereby, this study aimed to evaluate if there is a specific predominance of T or B lymphocytes and degranulated mast cells in the intestinal wall from dogs with IBD when compared with healthy controls.

## Materials and methods

### Dogs and Tissue Samples

Archived formalin-fixed paraffin-embedded (FFPE) intestinal biopsy pathology samples from dogs with a clinical and histopathological diagnosis of IBD were used for the study. In addition, samples from ten healthy dogs, with no history of antibiotic or immunosuppressive treatments two weeks before biopsy were included as controls. Only biopsy samples including both mucosa and submucosa, or the entire intestinal wall were considered in this study. Clinical inclusion criteria for these cases were vomiting, diarrhea, anorexia, weight loss, or some combination of these signs for at least 3 weeks, with no recent history of administration of immunosuppressive drugs or antibiotics. The histopathological inclusion criteria considered lymphocytic plasmacytic enteritis or colitis cases scored as marked, according to the histopathologic guidelines for the evaluation of gastrointestinal inflammation in companion animals recommended previously (42). Health status was assessed considering normal physical examination, and blood test results. Informed consent was obtained from all owners, and the study protocol was approved by the Ethics Committee of the Faculty of Life Sciences, Andres Bello University, Santiago, Chile. Endoscopic biopsy samples were collected for diagnostic and research purposes as described previously (43). All tissue samples were immediately placed in 10% neutral buffered formalin, processed conventionally and embedded in paraffin wax.

### Histology and Immunohistochemistry

All biopsies were cut (3μm) thick and stained with hematoxylin and eosin (H&E), toluidine blue, to detect granulated mast cells, and Masson’s trichrome staining, to assess fibrosis in lesions. Serial sections from samples were subjected to immunohistochemistry specific for T cells (CD3), B cells (CD79a) and mast cells (CD117). Briefly, after dewaxing, tissue sections were immersed in Tris-buffered saline (TBS 1X, pH 7.4). For antigen retrieval, slides were incubated for 20 minutes in citrate buffer (pH6.0) for CD3 and CD79a, or 15 minutes in proteinase K for CD117. Endogenous peroxidase was quenched with 3% hydrogen peroxide for 15 minutes. Non-specific binding was blocked by incubating slides for 20 minutes with a 2.5% normal horse blocking serum. Slides were incubated with the following primary antibodies: CD3 diluted 1:50 for 30 minutes, CD79a diluted 1:300 for 40 minutes, or CD117 diluted 1:300 for 20 minutes. Then, all slides were incubated with ImmPRESS™ HRP Reagent Kit Universal anti-mouse/rabbit IgG (Vector Laboratories, Burlingame, CA, USA) for 30 minutes. Primary antibody reactivity was detected by ImmPACT™ DAB Peroxidase Substrate Kit (Vector Laboratories, Burlingame, CA, USA) according to the manufacturer’s instructions, and all slides were counterstained with Mayer’s hematoxylin. Table 1 shows a summary of the primary antibodies used in the study.

**Table 1.**
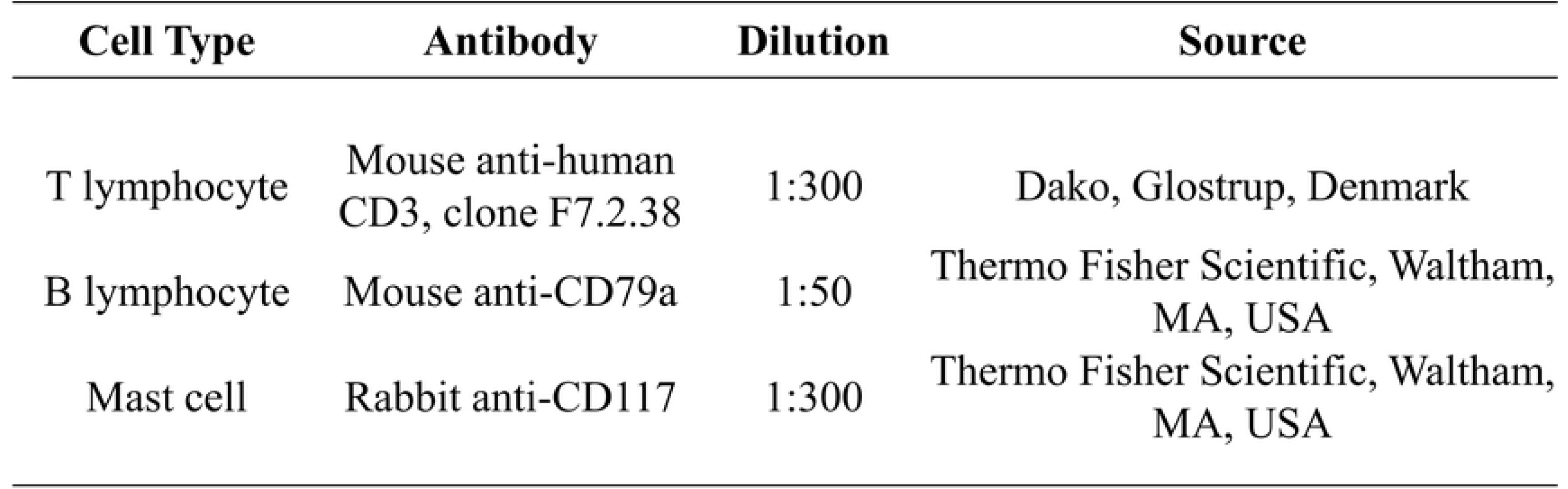
Primary antibodies used in the study.

For mast cell chymase and tryptase detection samples were incubated with MastCell Tryptase Abcam Rabbit monoclonal Clone ERP8476, and MastCell ChymaseThermo Mouse monoclonal Clone CC1. For double immune staining Polink TS-MMR-Hu A Kit, Polymer-HRP &AP triple staining kit was used.

### Examination of sections

Histological grading was assessed in H&E stained sections of intestine of dogs with IBD (n=39), according to WSAVA Gastrointestinal Standardization Group guidelines (42,44), by an experienced pathologist (CG). Vascular, degenerative and inflammatory histologic parameters were scored based on a scale from 0 to 3 as follows: normal (0), mild (1), moderate (2) and marked (3). Then, each parameter was assigned an importance factor (45,46) of 1 to 3 according to its histopathological relevance. We gave the maximal score to lacteal dilation, intraepithelial lymphocyte (IEL) and lamina propria lymphocyte infiltration compared to other features like fibrosis, villous stunting, epithelial injury and crypt distension. Also the height and width in both villi and lacteals was evaluated to assess villous stunting and crypt distention, as mentioned before (47). The assigned importance factor was multiplied by the score obtained in the first evaluation. Finally, we calculated the total corrected score by arithmetic means and classified cases into different severity groups as: normal (≤25% of the total score), mild (25–50% TS), moderate (50–75% TS) and marked (>75% TS).

Samples were evaluated with an Olympus microscope FSX-100 (Olympus, Center Valley, PA, USA) and then computer-assisted morphometric analyzed with the ImageProPlus TM 4.5 software (Media Cybernetics, Silver Springs, MD, USA). For scoring of intestinal CD3+ T lymphocytes, CD79a+ B lymphocytes, mast cells stained with blue toluidine and CD117+ mast cells, the number of cells in the lamina propria of ten appropriate fields (magnification at 40X) were quantified and arithmetic means were calculated for each. Results were expressed as the average of positive cells per high power field (HPF). Regarding for blue toluidine staining, we only considered granulated mast cells, those with visible metachromatic granules. Also morphologic features were taken into account that indicated that these cells were indeed mature granulated mast cells as described before (30).

### Statistics

The results were subjected to SPSS Statistics 22 software (SPPS Inc., Chicago, IL, USA) for analysis. Cell counts were assessed for normal distribution and non-parametric tests were performed. Differences between cell population numbers in dogs with IBD and controls were analyzed using the Wilcoxon test, and *P*≤0.05 was considered significant.

## Results

44,497 archived reports corresponding to all the tissues samples of dogs received in a 10 years period (2008-2017), by a Diagnostic Veterinary Laboratory (Citovet-Chile), were reviewed. Only 215 samples (0.48%) that showed histopathological findings compatible with IBD. Seventy-seven (88.5%) samples corresponded to a lymphocytic-plasmacytic enteritis, 92 (71.9%) samples to lymphocytic-plasmacytic colitis, 4 (4.6%) samples to eosinophilic enteritis, 5 (3.9%) samples to eosinophilic colitis and the remaining 37 samples corresponded to cases classified as non-specific enteritis or colitis. Distribution of the IBD cases from this study, is shown in Table 2.

**Table 2.** Distribution of anatomic regions affected and histological types of IBD cases.

| **Histopathological diagnosis** | **Enteritis (n=87)** | **Colitis (n=128)** |
| --- | --- | --- |
| IBD (n =215) |  |  |
| Lymphocytic plasmacytic | 77 (88.5%) | 92 (71.9%) |
| Eosinophilic | 4 (4.6%) | 5 (3.9%) |
| Non-specific | 6 (6.9%) | 31 (24,2%) |

A summary of the distribution of frequency of sex, age (Fig 1) and breed of the dogs diagnosed with IBD during the 10 years period, is shown in Table 3. Regarding healthy control dogs, 7 of them were females, and 3 males, age range between 1 to 6 years and more than half were crossbreed dogs.

**Table 3.**
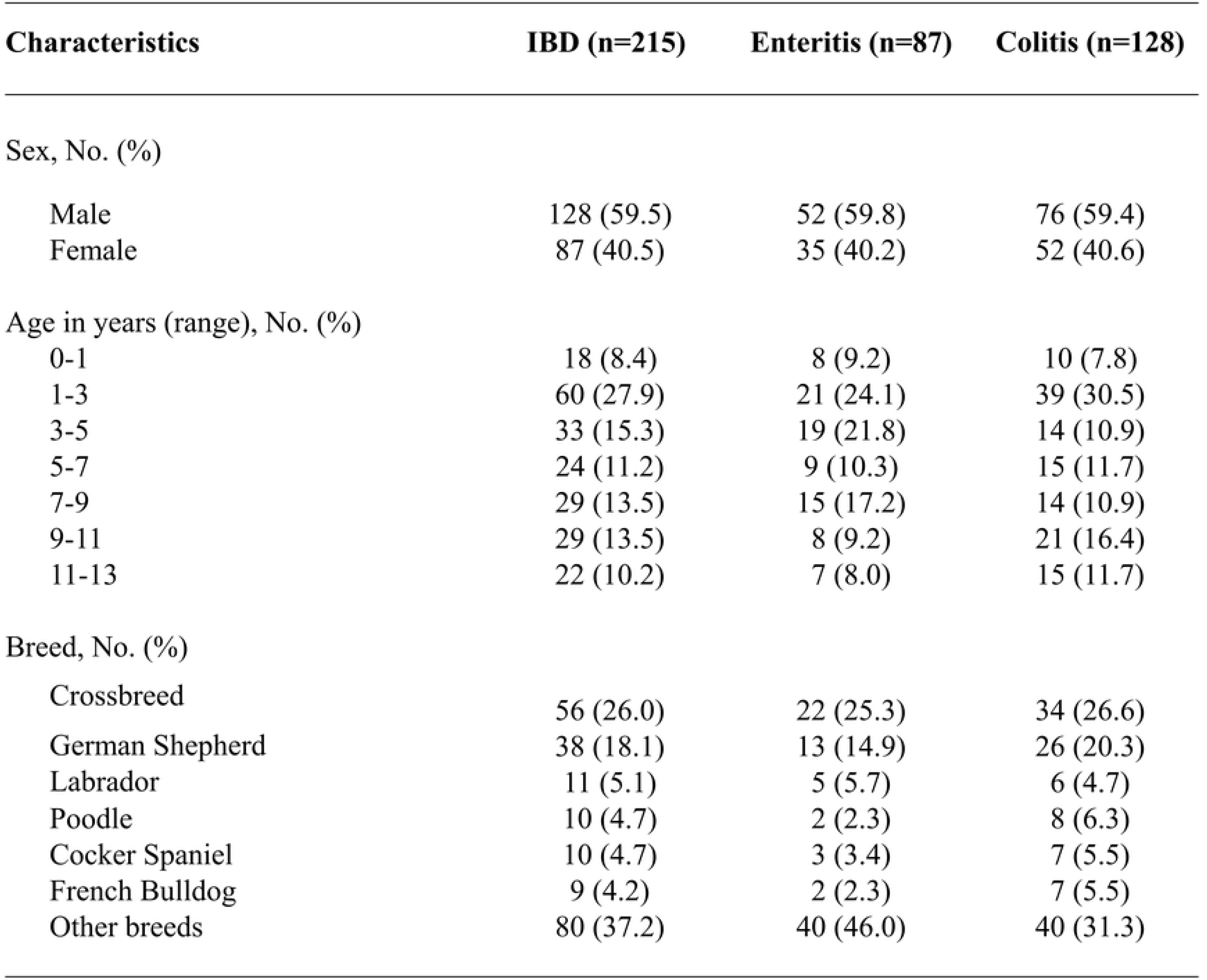
Distribution of frequency of sex, age and breed of dogs with IBD.

| Characteristics | IBD (n=215) | Enteritis (n=87) | Colitis (n=128) |
| --- | --- | --- | --- |
| Sex, No. (%) |  |  |  |
| Male | 128 (59.5) | 52 (59.8) | 76 (59.4) |
| Female | 87 (40.5) | 35 (40.2) | 52 (40.6) |
| Age in years (range), No. (%) |  |  |  |
| 0-1 | 18 (8.4) | 8 (9.2) | 10 (7.8) |
| 1-3 | 60 (27.9) | 21 (24.1) | 39 (30.5) |
| 3-5 | 33 (15.3) | 19 (21.8) | 14 (10.9) |
| 5-7 | 24 (11.2) | 9 (10.3) | 15 (11.7) |
| 7-9 | 29 (13.5) | 15 (17.2) | 14 (10.9) |
| 9-11 | 29 (13.5) | 8 (9.2) | 21 (16.4) |
| 11-13 | 22 (10.2) | 7 (8.0) | 15 (11.7) |
| Breed, No. (%) |  |  |  |
| Crossbreed | 56 (26.0) | 22 (25.3) | 34 (26.6) |
| German Shepherd | 38 (18.1) | 13 (14.9) | 26 (20.3) |
| Labrador | 11 (5.1) | 5 (5.7) | 6 (4.7) |
| Poodle | 10 (4.7) | 2 (2.3) | 8 (6.3) |
| Cocker Spaniel | 10 (4.7) | 3 (3.4) | 7 (5.5) |
| French Bulldog | 9 (4.2) | 2 (2.3) | 7 (5.5) |
| Other breeds | 80 (37.2) | 40 (46.0) | 40 (31.3) |

**Figure 1.**
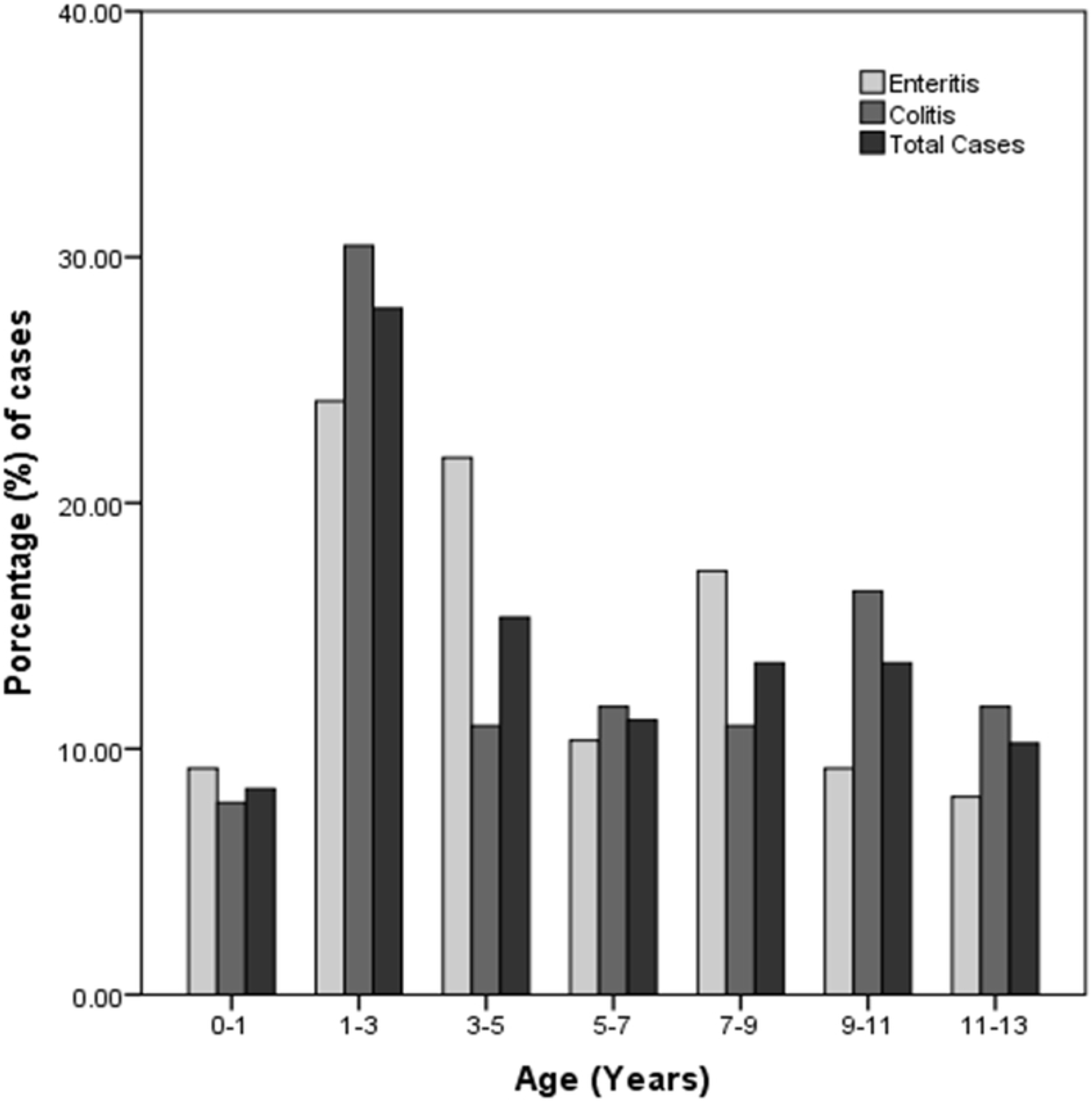
Age distribution of 215 dogs suffering from IBD.

Out of the 77 cases of dogs with a clinical and histopathological diagnosis of IBD lymphocytic-plasmacytic enteritis or colitis, that met the criteria established previously, thirty-nine FFPE intestinal samples, suitable for immunohistochemical study were selected for this study. Twenty-seven of thirty-nine intestinal samples (69.2%) corresponded to lymphocytic-plasmacytic enteritis (LPE). Nine of them, were classified as moderate LPE, and eighteen as marked LPE. Most of the marked cases showed significantly villous stunting, epithelial injury, lacteal distention, loss and dilation of crypts and a severe lymphocyte and plasma cells infiltration (Fig 3A).

The remaining twelve samples (30.8%) corresponded to lymphocytic-plasmacytic colitis (LPC). Ten of them were classified as moderate LPC, and the other two as marked LPC. Marked cases commonly showed villous stunting, dilatation, and loss of crypts, fibrosis, and a severe lymphocyte infiltration. Intestinal samples from all control dogs showed a normal histology.

### T and B lymphocyte populations

CD3+ T lymphocytes and CD79a+ B lymphocytes were detected in all cases. The median number of the different type of cells populations in IBD and control dogs are shown in Table 4.

**Table 4.** Comparison of mean lymphocyte and mast cells counts in dogs with IBD and control dogs.

| Cell type | Category | IBD group<br>(n=39) | Control group<br>(n=10) | P value |
| --- | --- | --- | --- | --- |
| T lymphocyte | CD3+ | 90.6 ± 77.20 | 40.0 ± 3.75 | <0.001 |
| B lymphocyte | CD79a+ | 28.5 ± 18.3 | 15.0 ± 5.0 | <0.001 |
| Mast cell | CD117+ | 12.0 ± 8.8 | 8.5 ± 3.28 | 0.013 |
| Mast cell | Toluidine Blue | 2.2 ± 7.5 | 7.95 ± 2.25 | 0.014 |

The total number of CD3+ T lymphocytes in the lamina propria of the intestine from dogs with IBD were significantly higher compared with control dogs (p<0.001). The number of CD3+ T lymphocytes of dogs with IBD was higher in the villi (Fig 2 and Table 5) and its number reduced toward the crypts in cases of LPE (Fig 3B), seeing a similar distribution in the colonic mucosa in cases of LPC. The total number of CD79a+ B lymphocytes in the lamina propria of dogs with IBD were significantly higher compared with control dogs (p<0.001). The distribution of CD79a+ B lymphocytes was similar in LPE and LPC cases, predominating near the crypt areas (Fig 2 and Table 5). The total number of CD3+ T lymphocytes was significantly higher (p<0.001) than the total number of CD79a+ B lymphocytes in the intestinal lamina propria of dogs with IBD. Regarding controls dogs, the total number of CD3+ T lymphocytes was significantly higher (p<0.001) compared with the total number of CD79a+ B lymphocytes.

**Table 5.** Distribution of the infiltrative lymphocyte and mast cells subpopulations in the intestinal wall mucosa in canine IBD and controls.

|  | LYMPHOCYTES |  |  |  | MAST CELLS |  |  |  |
| --- | --- | --- | --- | --- | --- | --- | --- | --- |
|  | CD3+ |  | CD79a+ |  | CD117+ |  | Toluidine Blue+ |  |
|  | IBD | CONTROL | IBD | CONTROL | IBD | CONTROL | IBD | CONTROL |
| **ZONE 1** | 3 | 2 | 1 | 2 | 2 | 2 | 1 | 1 |
| **ZONE 2** | 1 | 2 | 3 | 2 | 2 | 2 | 3 | 2 |
| **ZONE 3** | 1 | 1 | 1 | 1 | 1 | 1 | 1 | 2 |

\*0 (0% infiltrative cells) ,1 (<25% infiltrative cells) , 2 (25-50% infiltrative cells), 3 (>50% infiltrative cells)

**Figure 2.**
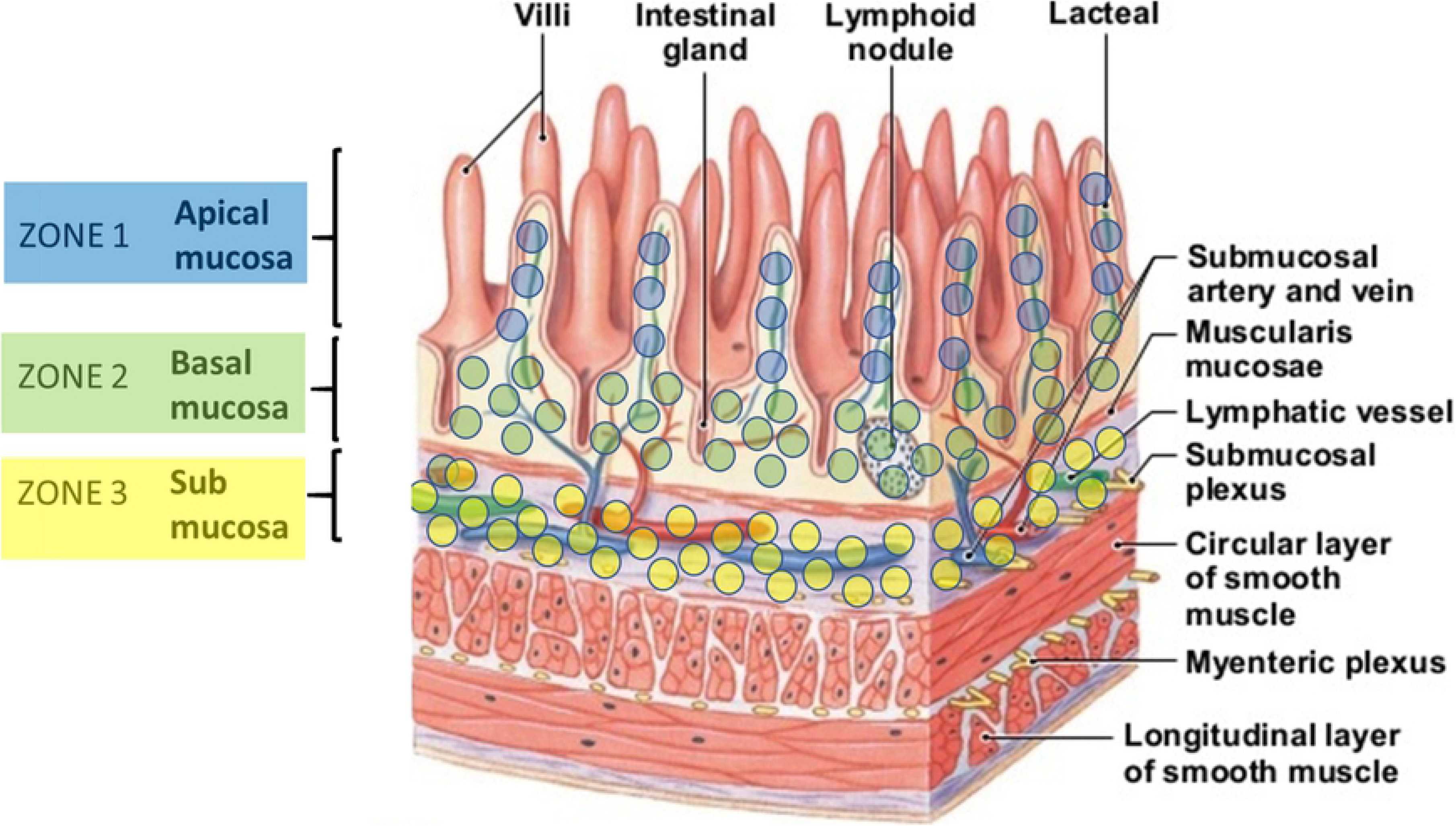
The histological organization of the intestinal wall and lymphocyte infiltration zones in canine IBD. Adapted from Martini et. al (2015) (48).

**Figure 3.**
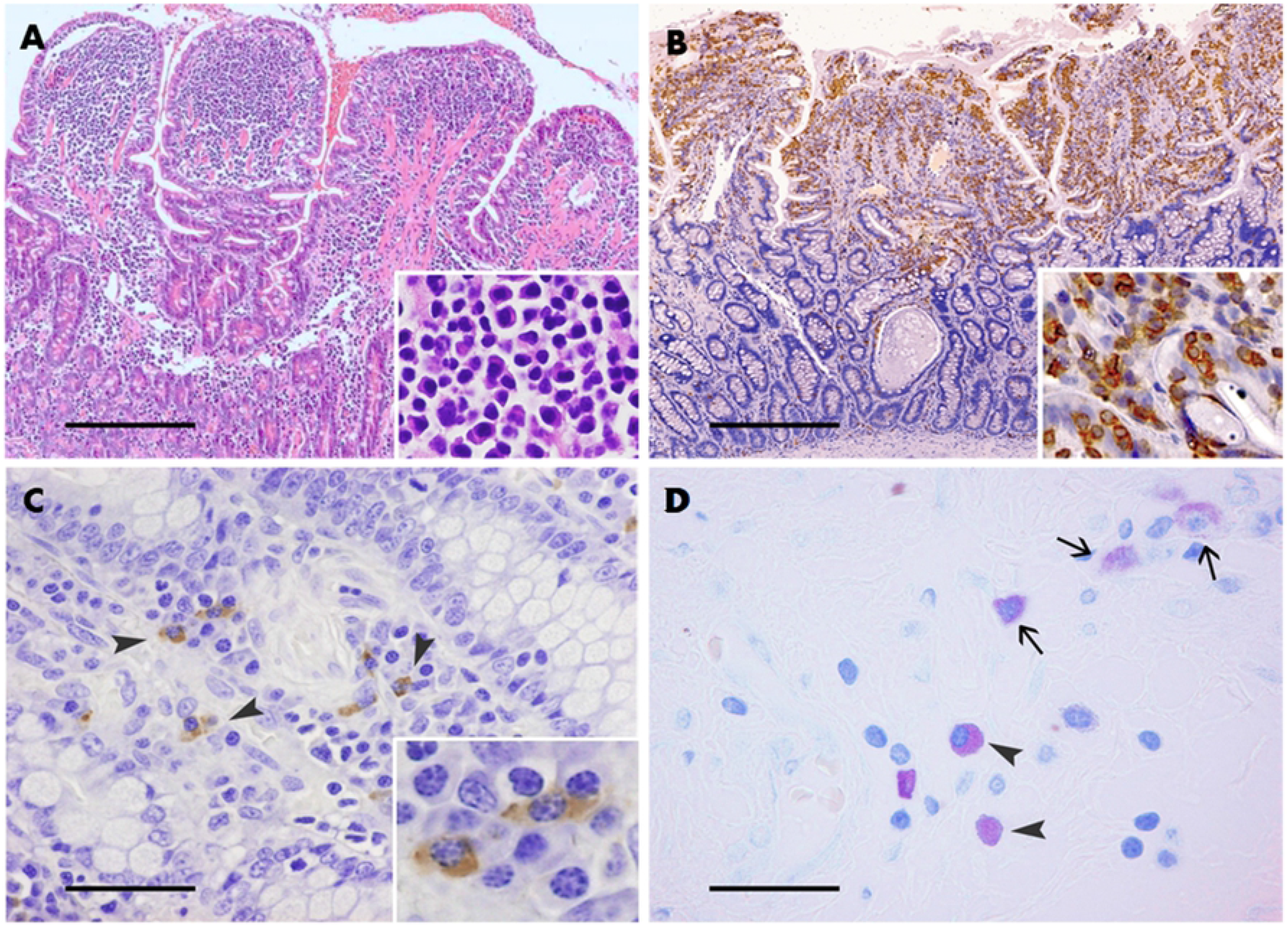
Lymphocytic-plasmacytic enteritis. A. Biopsy from a dog with lymphoplasmacytic enteritis of marked severity. There is an increase of lymphocytes, plasma cells and some macrophages in the lamina propria, marked villus stunting and fusion. Hematoxylin and eosin stain. Bar 300 μm. B. Large amount of CD3+ lymphocytes (brown color) are detected throughout the lamina propria of the villi using immunohistochemistry. Bar: 300 μm. C. CD117+ mast cells (arrowheads) scattered throughout the lamina propria of the villi using immunohistochemistry. Bar: 50 μm. D. Granulated mast cells (arrowheads) and degranulated mast cells (arrows) scattered throughout the lamina propria of the villi. Blue toluidine stain. Bar: 60 μm.

### CD117+ and blue toluidine mast cells

Mast cells were detected in all samples by both blue toluidine staining and CD117 immunophenotyping (Fig 3C and D). The total number of CD117+ mast cells in the lamina propria of the intestine from dogs with IBD was significantly higher compared with control dogs (p<0.013). However, the number of granulated mast cells detected with blue toluidine in samples from dogs with IBD was lower compared with control dogs (p<0.014). The total number of CD117+ mast cells detected in the lamina propria of dogs with IBD was significantly higher compared to the number of mast cells detected with blue toluidine in IBD dogs. (p<0.001). The distribution of mast cells was similar in LPE and LPC cases, located between the villous and close to the crypts (Fig 2 and Table 5).

### Chymase and tryptase detection in mast cells

Mast cells showed chymase and tryptase cytoplasmic expression in intestinal lamina propria. Some mast cells were only tryptase positive both in single stained samples (Fig 4A) or double stained samples (Fig 4C and E). Also, some mast cells were only chymase positive both in single stained samples (Fig 4B) or double stained samples (Fig 4C and F). On the other hand, there were mast cell were tryptase and chymase double positive (Fig 4D, E and F). The number of mast cells positive only to tryptase was significantly higher (p<0.001) than both mast cells positive only to chymase and mast cells double positive for tryptase/chymase. The double positive cells were significantly lower in number (p<0.001) to single positive cells to either tryptase o chymase.

**Figure 4.**
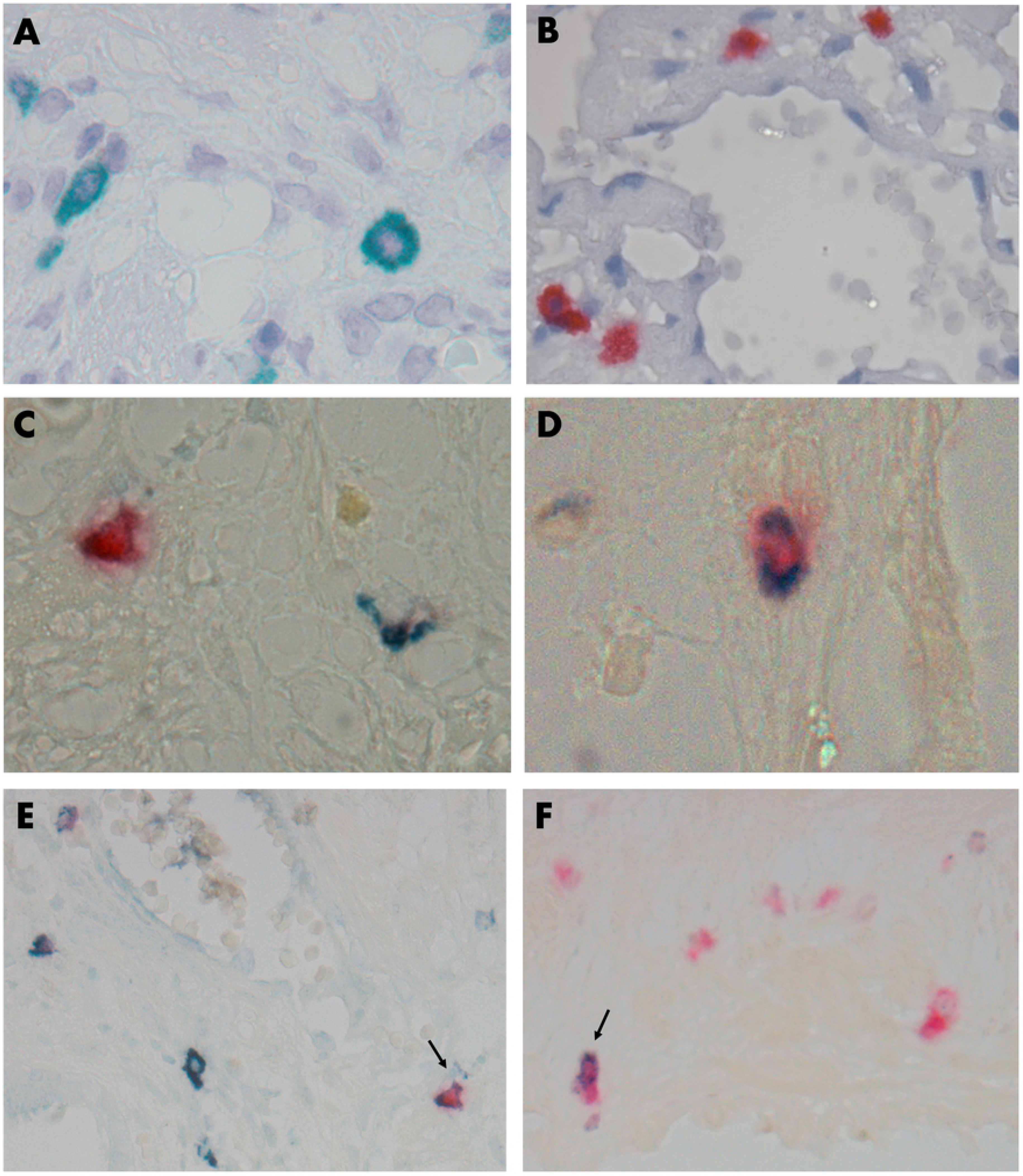
Positive reaction to chymase or tryptase expression or both in mast cells in canine small bowel lamina propria. Single immune staining for tryptase can be seen in Fig. A (green color) and for chymase in Fig. B (red color). Figure C to F show double immune staining for tryptase/chymase. Figure C shows mast cells positive to either tryptase (green color) o chymase (red color). Figure D shows a mast cell positive to both tryptase/chymase (green/red color). Figure E shows a group of mast cells stained for tryptase (green color) and one double stained (arrow). Figure F shows a group of mast cells stained for chymase (red color) and one double stained (arrow). Figs A to D 1000x. Figs E and F 400x.

## Discussion

In this study, different types of leukocytes were examined extensively in the intestinal layer of dogs with inflammatory bowel disease. After a retrospective study analysis of 44,497 FFPE samples of dogs, from 2010 to 2017, 215 samples presented histopathological findings compatible with IBD, giving a prevalence of 0,48% for this disease. Low IBD prevalence (1,99%) has also been reported Kathrani and colleagues, who found 546 patients diagnosed with IBD, after looking at the clinical records of 27463 patients treated at the Queen Mother Hospital for Animals, from the Royal Veterinary College during a six-year period (6). Although prevalence is similar in both studies, results are not necessarily comparable given the fact that our study considered only histopathological samples. German Shepherd dogs and crossbreeds were the most commonly affected breeds in our study. Similar results had also been reported in the United Kingdom by several studies (6,49–51), suggesting that German shepherd dogs are at increased risk of developing IBD. Authors identified several candidate genes by genome-wide association studies, showing a genetic predisposition of this breed. However, our study did not include any genetic analysis, and results may support this hypothesis, but could also be reflecting a sample bias (See Table 3). No significant difference regarding gender or age was found in the IBD group, as similarly reported by other studies (6,52).

Unexpected difficulties were experienced when processing the endoscopic biopsy samples, particularly related to a low amount of obtained material, localization of the lesions and improper orientation of the tissue. Willard and colleagues and Hall reported that the quality and amount of endoscopically obtained tissue samples has a profound effect on their sensitivity for identifying certain lesions (43,53). In addition, almost all endoscopic biopsies were sampled from duodenum or/and colon, and few of them had ileum. This may be relevant as Casamian-Sorrosal and colleagues found poor agreement between histopathological findings from duodenal versus ileum biopsies with abnormalities sometimes being more readily detected in the ileum (54), thus the collection of concurrent duodenal and ileum endoscopic biopsies is recommended (55). As we were not able to control how the samples were collected and processed, only those showing lesions consistent with IBD were included in this study.

Most of IBD cases corresponded to lymphocytic-plasmacytic enteritis/colitis. Previous studies demonstrated that lymphocytic-plasmacytic enteritis/colitis is the most common form of IBD in dogs (1,56,57). Many researchers follow the endoscopic, biopsy, and histopathologic guidelines for the evaluation of gastrointestinal inflammation in companion animals (42,44) to assess the pathological lesions found in chronic gastroenteropathies in dogs with IBD. Villus atrophy, lacteal dilatation, and the presence of inflammatory infiltrate are evaluated and graded with this system, however, these guidelines do not rank the lesion according to how important it is to the organ function. Different studies proposed the use of a standardized method that allows the quantification of organ damage, extent and pathological importance of changes in a fish species (45,46,58). This standardized tool for the assessment of histological lesions can be applied to different organs. Hereby, a similar approach was put forward to classify IBD scoring in dogs. Using the proposed method, it was possible to categorize marked and moderate cases regarding the traditional guideline that classifies them initially as moderate and mild respectively.

Most LPE cases were classified as marked, while most LPC cases were scored as moderate. These findings might be related with the fact that most patients are referred for endoscopy and histopathological examination only when clinical signs have persisted for a long period of time or when patients do not respond to conventional treatment. In humans, diagnosis of IBD is often delayed, particularly in Crohn’s disease (CD) due to the variety of its clinical signs, being opposed to ulcerative colitis (UC), where symptoms appear earlier (59). Further, a long diagnostic delay was associated with a poor outcome when IBD-related surgery was performed in CD and UC patients (60). No recent studies regarding how a diagnosis delay affects the severity of IBD in dogs were found, however, it is possible that a similar scenario is seen, being moderate cases detected faster in lymphocytic-plasmacytic colitis than in lymphocytic-plasmacytic enteritis due to earlier signs manifestation. More studies are necessary to elucidate this issue.

A significantly higher number of CD3+ T lymphocytes, CD79a+ B lymphocytes, and CD117+ mast cells were detected in the lamina propria of dogs with IBD compared with healthy dogs. Changes in the number of CD3+ lymphocytes were similar to those found in other studies (61,62) in which CD3+ T cells were higher in the lamina propria of dogs with IBD. These changes in the lymphocytic composition could be related to a variety of components affecting the immune system of those individuals (5). Jergens found that a resistance to apoptosis by T cells could also contribute to the typical cell accumulation in IBD dogs (63). A higher number of T cells in the lamina propria of the gut compared with B cells was also found, which may be related to the cytokine profile present in this disease (16,64,65), facilitating the accumulation of T cells in the lamina propria. CD117+ mast cells were detected in a higher number in the lamina propria of the intestine from dogs with IBD similarly in other study (66), and also observed in intestine samples from cats with IBD (67).

Conversely, a decreasing number of metachromatically stained mast cells was reported (15,68), suggesting reduced lamina propria mast cell numbers in dogs with IBD. This apparently occurs because the staining of mast cells by immunohistochemistry, but not by toluidine blue, is independent of whether mast cells are degranulated or not. Our results suggest that the number of mast cells in the lamina propria of intestine from dogs with IBD does not necessarily increase significantly, but the portion of degranulated mast cells is higher. These findings point out that whether mast cells are increased in number or not they become activated and degranulated during IBD in dogs. Mast cells degranulation will also trigger the recruitment of more inflammatory cells, like T lymphocytes, in the intestinal layer (26,36), intensifying the typical lymphocyte infiltrative pattern seen in this disease.

In this study the number of mast cells positive only to tryptase was significantly higher (p<0.001) than both mast cells positive only to chymase and mast cells double positive for tryptase/chymase. Tryptase is a potent growth factor for epithelial cells, smooth muscle and fibroblasts, it also regulates gastrointestinal smooth muscle activity and intestinal transport (69). It also develops as an anticoagulant, fibrinolytic agent, activator of PAR-2 (Protease-Activated Receptor) (activated by trypsin and mast cell tryptase, associated with physiological processes aggregation, cell proliferation, and in contraction or relaxation of the smooth muscle) (70), improvement of vascular permeability, angiogenesis, inflammation and hyperactivity of the smooth muscle of the airways (71). Whereas chymase helps maintain the function of the intestinal barrier (37), participates in defense against parasites (72), promotes epithelial permeability, and maintains blood pressure during anaphylaxis by the generation of angiotensin II, Chymase and Cathepsin G destroy several cytokines associated with inflammation, which causes the decrease or end of inflammation (37).

In our study CD3+ T leucocytes were predominantly located in the villous region, however, CD79a+ B cells and CD117+ mast cells were located close to crypt region, very similar to other studies (61,73). There is no clear reason for this distribution, but it is possible that T lymphocytes are higher in villi because of a higher presentation of antigen associated to the molecule histocompatibility complex by mononuclear cells (74), and this could be related to the close contact of stimulant bacteria in the villi and the crypts. In mice, mast cells reside in the muscle/submucosa region (75) and after they are activated by substance P produced by lymphocytes and macrophages (76), they move into the lamina propria and the epithelium where inflammatory mediators such as serotonin and proteases are released (25). Mast cells located around the crypts could be associated with close contact of environmental antigens, because they recognize molecular motifs (pattern recognition molecules) which trigger a subsequent immune response attracting lymphocytes to the site of the lesion (77,78). Thus, a clear and strong relation between mast cells and T lymphocytes could possibly explain the changes observed in the intestinal layer of dogs affected with IBD.

In summary, our results show that CD3+ lymphocytes and CD117+ mast cells were the predominant leukocyte population in the intestine of dogs with IBD compared to control dogs and found that CD3+ lymphocytes mainly occupy villous tips while CD79a+ lymphocyte and CD117+ are distributed close to the crypts. Changes in the number of T lymphocytes and mast cells may be considered as an additional criteria tool for the diagnosis of IBD in dogs, as increased numbers of both cell types were found as an indicator of disease severity. Also, the number of activated mast cells detected by CD117 immunophenotyping was higher than the number of granulated mast cells detected by blue toluidine staining, showing that degranulation is probably occurring in this disease. The information generated from this study suggest an important role of both T lymphocytes and mast cells in the immunopathology of IBD in dogs. The specific role of these leukocytes populations in IBD onset and development should be further investigated to allow the development of medical treatments focused on their regulation. Although diagnostic guidelines for IBD report the parameters for mild IBD scoring, in our experience, differences between structural and infiltrative changes between normal cases and mild IBD are difficult to demonstrate, and misclassification may occur. Considering this, only moderate and severe cases were included in this study. However, according to the results obtained in his study differences in both infiltration by lymphocyte subpopulations and their tissue location as well as estimation of mast cell number should be also investigated in mild cases.

## Acknowledgments

The author acknowledges funding provided by FONDECYT Regular 1121202, Universidad Andres Bello, Santiago, Chile. We thank Ivan Contreras for assistance in protocol and experiment design.

## Conflict of Interest Statement

All the authors declare that they do not have any conflicts of interest respect to the research, authorship and/or publication of this article.

